# T cell immunity does not age in a long-lived rodent species

**DOI:** 10.1101/259374

**Authors:** M. Izraelson, T.O. Nakonechnaya, A.N. Davydov, M.A. Dronina, D.A. Miskevich, I.Z. Mamedov, L.N. Barbashova, M. Shugay, D.A. Bolotin, D.B. Staroverov, E.Y. Kondratyuk, E.A. Bogdanova, S. Lukyanov, I. Shams, O.V. Britanova, D.M. Chudakov

## Abstract

Numerous studies have demonstrated that the percentage of naïve T cells and diversity of T cell receptor (TCR) repertoire decrease with age, with some findings likewise suggesting that increased repertoire diversity may be associated with longer lifespan and healthy aging. In this work, we have analyzed peripheral TCR diversity from humans, mice, and blind mole-rats (*Spalax spp*.)—long-lived, hypoxia- and cancer-tolerant rodents. We employed a quantitative approach to TCR repertoire profiling based on 5’RACE with unique molecular identifiers (UMI) to achieve accurate comparison of repertoire diversity, which also required development of specific wet lab protocol and TCR gene reference for *Spalax*. Our direct comparison reveals a striking phenomenon. Whereas TCR diversity of mice and humans decreases with age, resulting primarily from the shrinkage of the naive T cell pool, *Spalax* TCR diversity remains stable even for the animals that reach extreme old age (15-17 years). This indicates that T cell immunity does not meaningfully age in long-lived rodents, at least in terms of the classical understanding of immunosenescence, which is associated with the accumulation of large numbers of memory clones. We suggest that the extraordinary longevity of *Spalax* may be attributable at least in part to the distinctive organization of their T cell immunity. Our findings should therefore encourage a close re-examination of the contribution of immunosenescence to life span in mammals.

---

The T cell receptor (TCR) diversity of naïve T lymphocytes represents a precious collection of keys from which antigen-specific variants are selected, conferring protection for the host against new challenges. This diversity allows the adaptive immune system to recognize variable and evolving pathogens, as well as neoantigens produced by cancer cells^1^. It is also important to provide diverse regulatory activities^2^ and thus to sustain the generally balanced state of adaptive immunity.

Naïve T cells are actively produced in young mice until they are approximately 3 months old, and in humans until they are approximately 20 years old. Along with accumulation of expanded memory T cell clones, counts of naïve T cells in peripheral blood decrease linearly with age, as reflected by a proportional decrease in observed TCR diversity^3-5^. This age-related effect on T cell immunity is one of the most prominent hallmarks of the whole aging process, leading to increased susceptibility to infections and cancer in the elderly^3, 4^. Notably, cytomegalovirus (CMV) infection enhances immunosenescence, and the absence of CMV is associated with an increased percentage of naïve CD4 T cells in the elderly and longer life expectancy in humans^3, 6^.

Here we raised the question of whether the exceptional longevity of certain rodent species such as the naked mole rat or blind mole-rat (*Spalax* spp., **Fig. 1**)—which live for 30 and 20 years, respectively^7^ versus 2–3 years for mice and rats, and which mortality does not increase with age^8^—could be associated with slower aging of T cell immunity. To verify our hypothesis, we performed TCR alpha (TRA) and beta (TRB) CDR3 repertoire profiling for spleen samples derived from 19 *Spalax* animals aged 0–17.5 years and 14 C57Bl/6 mice aged 3–24 months. We employed cDNA 5’-RACE protocol with unique molecular identifiers (UMI)^5^, using primers specifically designed for mice and *Spalax* (**Table 1**). UMIs drastically improve quantification and thus allow accurate comparison of TCR diversity between samples^5, 9, 10^. This protocol also takes advantage of universal primer sites located in the constant regions of the TRA and TRB, and is therefore independent of the particular sequences of the diverse TRBV gene segments, previously unknown for *Spalax*. TCR libraries were subjected to 150+150-nt paired-end sequencing on an Illumina HiSeq. UMI-based data TCR repertoire analysis was performed using MiGEC, MiXCR, and VDJtools software.

**Figure 1.**
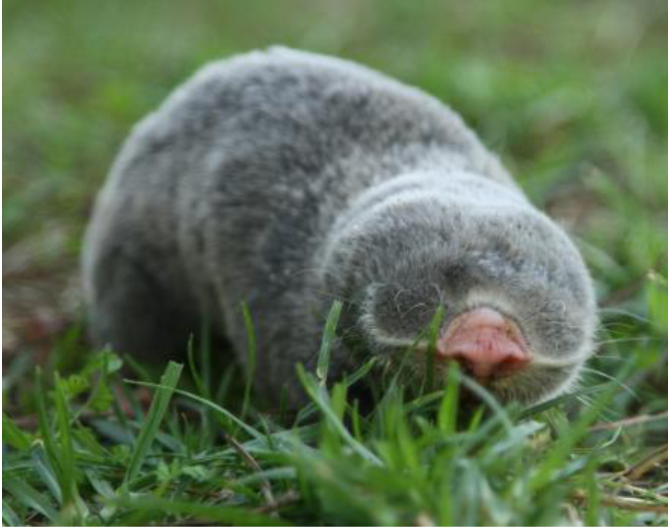
Spalax.

In striking contrast to mice and humans, we found that peripheral TCR diversity of *Spalax,* albeit moderate, is highly stable and does not decrease with age (**Fig 2**). This observation indicates that long-lived rodents possess distinct mechanisms for sustaining efficient T cell immunity, where clonal memory expansions are not occupying significant volume and naïve cells represent the majority of the peripheral T cell population even at very advanced ages. This could be explained by prolonged thymic activity or increased production of T cell progenitors by haematopoietic stem cells, but is most probably attributable to more rapid turnover of memory T cells. Lack of large clonal expansions at any age suggests that *Spalax* may have relatively poor long-term T cell memory. This would leave the animals well-prepared to efficiently withstand new challenges due to the stably high diversity of TCR repertoire, albeit at the cost of diminished enhancement of responses to previously encountered pathogens.

**Figure 2.**
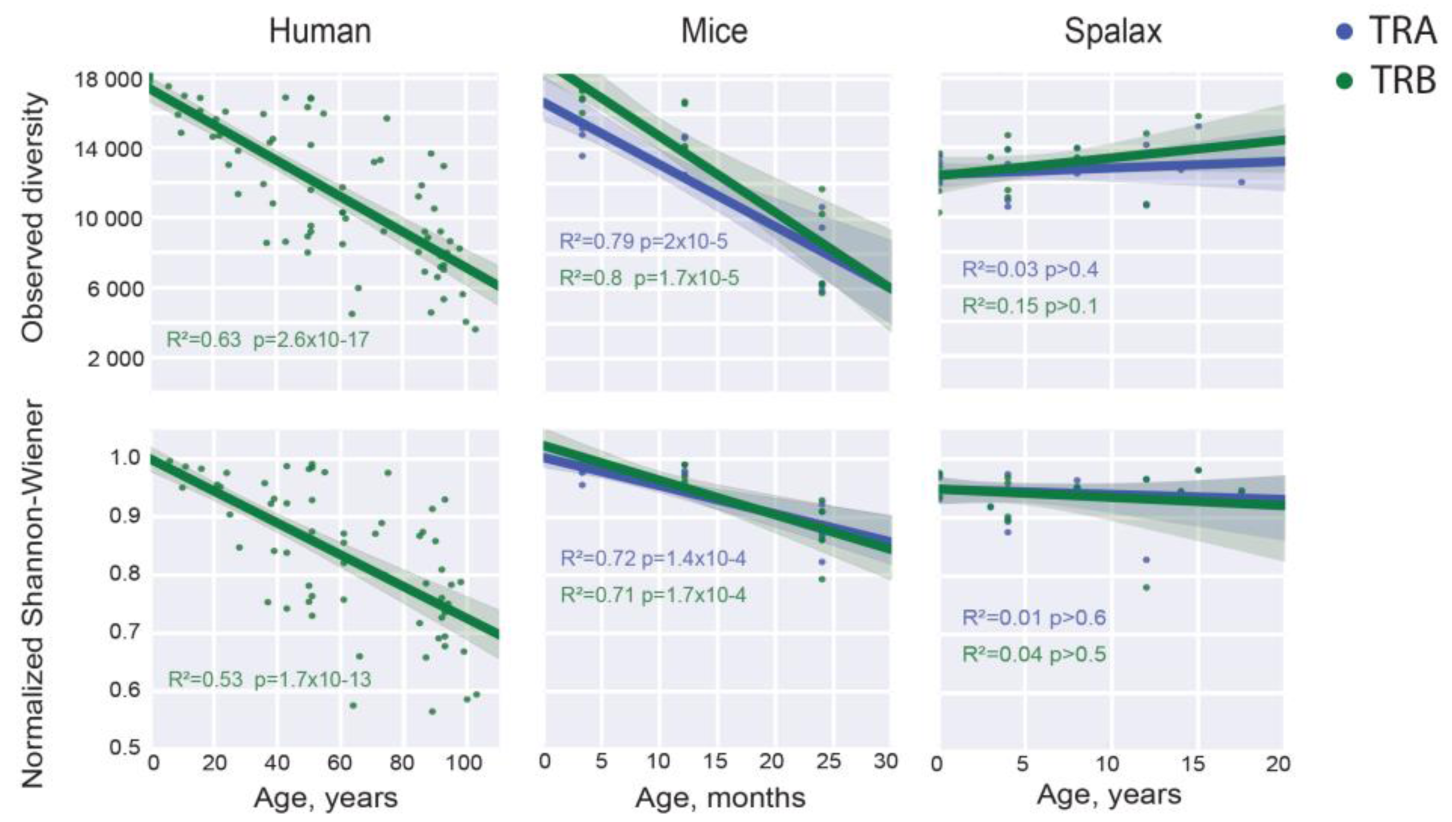
Diversity of T cell receptors and age. TRA and TRB in-frame nucleotide CDR3 diversity for human peripheral blood, C57Bl/6 mouse spleen, and *Spalax* spleen samples from subjects of different ages. Diversity was estimated in terms of the directly observed number of variants (top) and the normalized Shannon-Wiener index (bottom) for the samples which were normalized to an equivalent number of 18,300 unique TCR cDNA molecules labeled with UMI. Human data was re-analyzed from Ref. ^5^. P-value is a two-sided p-value, where the null hypothesis is that the slope is zero, using a Wald Test with t-distribution of the test statistic.

This discovery raises intriguing questions concerning possible strategies of adaptive immunity organization, and should initiate a new wave of studies into the relationship between aging and immunity. Alongside the intense investigations of genomics and metabolomics in long-lived rodents^11, 12^, this new avenue of immunological research should lead us to a much more fundamental and holistic understanding of the aging process in mammals.

## Acknowledgments

The work was supported by the Russian Science Foundation project ? 16-15-00149.

## Materials and Methods

### Animals

Blind mole-rats (hereafter, *Spalax*) were captured in the field and individually housed in cages (L 45 cm x W 30 cm x H 20 cm) with sawdust bedding. The animals were maintained at 22°C, and fed with fresh vegetables. Ages of the animals were estimated as follows. For adult animals, two years were added to the period of time spent in housing up until the experiment. Therefore, the ages of the adult animals represent minimum estimates. Newborn individuals were taken from underground nests, and their age was estimated according to size and morphological parameters. Animals were sacrificed with an overdose of an anaesthesia agent, followed by immediate splenectomy. Tissues were then flash-frozen in liquid nitrogen and stored at -80°C for further analyses. *Spalax* work was accomplished under the supervision of the University of Haifa Ethics Committee, approval #s 316/14 and 420/16.

C57BL/6J mice were purchased from the Puschino Animal Breeding Center (Moscow, Russia) and housed in the SPF facility of the Institute of Bioorganic Chemistry of the Russian Academy of Sciences (IBCH RAS). The work was accomplished under the supervision of the IBCH IACUC and following the Regulations of the Ministry of Health of the Russian Federation. All procedures were approved by IBCH IACUC protocol No.168.

### Tissue collection and RNA isolation

C57BL/6J mice spleens were isolated and chopped in PBS. White mononuclear cells were extracted using Ficoll-Paque (Paneco, Russia) density-gradient centrifugation, put in RLT buffer (Qiagen) and frozen at -70°C. RNA was extracted using the RNeasy Micro Kit (Qiagen). Frozen pieces of the *Spalax* spleens were homogenized using Trizol (Invitrogen) in liquid nitrogen, and RNA was isolated according to the manufacturer’s manual. Total RNA was dissolved in 30 µl of sterile water. RNA concentration and quality were evaluated with a Qubit 3.0 Fluorometer (Invitrogen) and gel electrophoresis analysis.

### Primer design for cDNA synthesis and target amplification of the *Spalax* TCR-alpha/beta libraries

Primers specific to the gene segments encoding the constant regions of the beta and alpha chains of *Spalax* TCR were designed according to the homology alignment analysis of corresponding mouse, human, and rat gene segments versus *Spalax* RNA-Seq data^13^. The nucleotide homology between *Spalax* and mouse constant gene segments of the TCR alpha and beta constant regions was more than 85%.

### cDNA synthesis

cDNA libraries were obtained using 5’-RACE^14^ with unique molecular identifiers (UMI) as described in Ref. ^15^, with minor modifications. cDNA synthesis was performed immediately after RNA extraction. We employed 3 µg of total RNA per 30 µl cDNA synthesis reaction for each sample from mice or *Spalax*. RNA was incubated with 0.5 mM reverse primers for TRBA and TRBC (see **Supplementary Table 1** for the oligonucleotides used) at 65°C for 2 min, then cooled to 42°C for primer annealing. Reverse transcription mix was then added, consisting of 1 mM DTT, 0.5 mM of each dNTP, 5 U/ml SMARTScribe reverse transcriptase, and 0.5 mM 5’-template switch adapter SmartNNNa, in SMARTScribe buffer (Clontech). This mixture was incubated for 60 min at 42°C for cDNA synthesis and template switching. To remove the deoxyuridine-containing template switch adapter, the cDNA was treated with uracil–DNA glycosylase (New England BioLabs, 5 U/ml) for 40 min at 37°C. Each cDNA synthesis reaction was purified with a MinElute PCR purification kit (Qiagen), with an elution volume of 30 μl.

### PCR amplification 1

First-strand cDNA was used for the 1st PCR with a gene-specific primer and step-out mix of Smart20 and Step1 primers. The longer primer, Smart20, has a 3’ segment that corresponds to the SmartNNNa adapter, and a 5’ segment that serves as a template for the Step1 primer. All cDNA was used in the first PCR. The 50 μl reaction mix included 10 μl of cDNA, 5X buffer for Q5 DNA polymerase (New England BioLabs), 0.2 mM of each dNTP, Q5 DNA polymerase (New England BioLabs), 0.02 mM Smart20 primer, 0.2 mM Step1 primer, and 0.2 mM reverse primers for TCR β- and α-chain amplification. TCR α- and β-chains were amplified together in the same round of PCR. The amplification program was as follows: 94°C for 2 min, then 20 cycles of 94°C for 10 s, 60°C for 10 s, and 72°C for 40 s. PCR products were purified using a QIAquick PCR purification kit (Qiagen) and eluted in 30 μl of elution buffer.

### PCR amplification 2

1 μl of the first PCR product was used as a template for the second PCR reaction. In order to combine PCR libraries from multiple samples in a single Illumina run, we introduced sample barcodes within the forward and reverse primers. The 50 μl reaction mix included buffer for Q5 DNA polymerase, 0.2 mM of each dNTP, 0.2 mM forward primer M1S, and a bcj or acj reverse primer for the TRBC and TRAC gene segments, respectively. The amplification program was as follows: 94°C for 2 min, then 9-10 cycles of 94°C for 10 s, 60°C for 10 s, 72°C for 40 s. The samples were combined together and purified using the QIAquick PCR purification kit.

### Sequencing

Illumina adapters were ligated, the libraries were amplified and then sequenced in a HiSeq4000 150+150 nt paired-end run, with 30% random DNA added. Additionally, we performed MiSeq 300+300 nt paired-end sequencing to generate long-range data that were used to build the complete *Spalax* TCR alpha and beta gene reference for MiXCR.

### Data analysis

Raw sequencing data were analyzed using MIGEC software^16^ to group sequencing reads carrying the same UMI and thus covering the same cDNA molecule. In-frame TCR alpha and beta CDR3 repertoires were extracted using MiXCR software^17^.

To efficiently extract the *Spalax* TCR repertoires, we developed a *Spalax* TCR gene reference (doi:10.5281/zenodo.834389; doi:10.5281/zenodo.1127674). To this end, we aligned 5’RACE TCR sequencing data to the *Spalax* reference genome assembly^18^ using BWA aligner, regions with coverage greater than 10 were extracted and inspected for the presence of specific features of each segments such as two exons, recombination signal sequences (RSS, mouse RSS available on www.imgt.org) for Variable genes, one exon and RSS for Diversity and Joint segments. Genomic coordinates of all genes were extracted and processed with repseqio software (https://github.com/repseqio/repseqio). The resulting extraction efficiency in terms of percentage of sequencing reads that contain identified TCR CDR3 reads was 95% and 74% for TRA and TRB genes respectively for mouse using mouse reference, and 94% and 82% for TRA and TRB genes respectively for *Spalax* using *Spalax* reference.

We applied VDJtools software^19^ for down-sampling to the same count of randomly-chosen 18300 unique UMI-labeled cDNA molecules (the size of the smallest dataset), and for comparative analysis of diversity metrics.

## References

1. Stronen, E. et al. Targeting of cancer neoantigens with donor-derived T cell receptor repertoires. Science 352, 1337–1341 (2016).

2. Li, M.O. & Rudensky, A.Y. T cell receptor signalling in the control of regulatory T cell differentiation and function. Nat Rev Immunol 16, 220–233 (2016).

3. Nikolich-Zugich, J. Aging of the T cell compartment in mice and humans: from no naive expectations to foggy memories. J Immunol 193, 2622–2629 (2014).

4. Qi, Q. et al. Diversity and clonal selection in the human T-cell repertoire. Proc Natl Acad Sci U S A 111, 13139–13144 (2014).

5. Britanova, O.V. et al. Dynamics of Individual T Cell Repertoires: From Cord Blood to Centenarians. J Immunol 196, 5005–5013 (2016).

6. Wertheimer, A.M. et al. Aging and cytomegalovirus infection differentially and jointly affect distinct circulating T cell subsets in humans. J Immunol 192, 2143–2155 (2014).

7. Tacutu, R. et al. Human Ageing Genomic Resources: new and updated databases. Nucleic Acids Res 46, D1083–D1090 (2018).

8. Ruby, J.G., Smith, M. & Buffenstein, R. Naked Mole-Rat mortality rates defy gompertzian laws by not increasing with age. eLife 7 (2018).

9. Kivioja, T. et al. Counting absolute numbers of molecules using unique molecular identifiers. Nat Methods 9, 72–74 (2012).

10. Izraelson, M. et al. Comparative analysis of murine T-cell receptor repertoires. Immunology 153, 133–144 (2018).

11. Kim, E.B. et al. Genome sequencing reveals insights into physiology and longevity of the naked mole rat. Nature 479, 223–227 (2011).

12. Lewis, K.N. et al. Unraveling the message: insights into comparative genomics of the naked mole-rat. Mammalian genome : official journal of the International Mammalian Genome Society 27, 259–278 (2016).

13. Malik, A. et al. Genome maintenance and bioenergetics of the long-lived hypoxia-tolerant and cancer-resistant blind mole rat, Spalax: a cross-species analysis of brain transcriptome. Scientific reports 6, 38624 (2016).

14. Matz, M. et al. Amplification of cDNA ends based on template-switching effect and step-out PCR. Nucleic Acids Res 27, 1558–1560 (1999).

15. Egorov, E.S. et al. Quantitative profiling of immune repertoires for minor lymphocyte counts using unique molecular identifiers. J Immunol 194, 6155–6163 (2015).

16. Shugay, M. et al. Towards error-free profiling of immune repertoires. Nat Methods 11, 653–655 (2014).

17. Bolotin, D.A. et al. MiXCR: software for comprehensive adaptive immunity profiling. Nat Methods 12, 380–381 (2015).

18. Fang, X. et al. Genome-wide adaptive complexes to underground stresses in blind mole rats Spalax. Nature communications 5, 3966 (2014).

19. Shugay, M. et al. VDJtools: Unifying Post-analysis of T Cell Receptor Repertoires. PLoS computational biology 11, e1004503 (2015).

